# TRPV1 activation relies on hydration/dehydration of nonpolar cavities

**DOI:** 10.1101/114546

**Authors:** Marina A. Kasimova, Aysenur Yazici, Yevgen Yudin, Daniele Granata, Michael L. Klein, Tibor Rohacs, Vincenzo Carnevale

## Abstract

TRPV1 promotes cationic currents across cellular membranes in response to multiple stimuli such as increased temperature, binding of chemicals, low pH and voltage. The molecular underpinnings of TRPV1 gating, in particular the mechanism of temperature sensitivity, are still largely unknown. Here, we used molecular simulations and electrophysiology to shed light on the closed to open transition. Specifically, we found that gating of TRPV1 relies on the motion of an evolutionarily conserved amino acid (N676) in the middle of the S6 helix. On rotation, the side chain of this asparagine faces either the central pore or the S4-S5 linker. Only in the former case is the central pore hydrated and thus conductive. Interestingly, when N676 rotates toward the linker, we observe hydration of four so far unreported small nonpolar cavities. Based on these findings, we propose a model for TRPV1 gating involving the dynamic hydration of these four cavities. Free energy calculations indicate that this gating mechanisms is markedly temperature dependent favoring the open state at high temperature. On the basis of this model, which is able to rationalize a wealth of seemingly conflicting and/or unrelated experimental observations, we predicted the behavior of two single residue mutants, M572A and F580Y, the consequences of which we confirmed experimentally.

## INTRODUCTION

TRPV1 is one of the most studied members of the TRP family of ion channels^1,2^. As many other members of the TRP family, TRPV1 shows multiple activation modalities, including activation by high temperatures, low extracellular pH, membrane depolarization and binding of vanilloids^1,3–14^. In animals, TRPV1 is mostly expressed in small- to medium-diameter neurons of primary sensory ganglia of a peripheral nervous system, where, upon activation, it transmits a painful signal through the dorsal root ganglia into the thalamus. Direct involvement of TRPV1 in nociception makes it an attractive target to silence pain pathways and hence one of the ion channels of pharmacological interest^15–17^.

Several recent cryo-EM studies have revealed the architecture of TRPV1^18–20^. This protein is a tetrameric assembly with a transmembrane (TM) domain and a large cytoplasmic one. The latter is composed of the N-terminal ankyrin repeats, the linker, the pre-S1 helix and the C-terminus domain. The TM domain resembles closely that of voltage-gated ion channels and is composed of two main moduli, the S1-S4 and the pore domain, which are connected through the S4-S5 linker. The pore domain creates a pathway for ions through the membrane and is composed of the S5 and S6 helices from all four subunits. Finally, TRPV1 has an amphipathic helix called the TRP box, which connects the S6 helix with the C-terminus domain.

Several structures of TRPV1 resolved in different conformational states provided significant insight into gating of this channel^18–20^. In particular, it was shown that TRPV1 has two gates, upper and lower, located, respectively, at the level of the selectivity filter and of the S6 bundle crossing. Both of these gates are open in the resiniferatoxin-bound (RTX-bound) state and closed in the apo state; in the capsaicin-bound (CAP-bound) state, the upper gate is closed, while the lower gate is partially dilated. A possible molecular mechanism for the transition between the apo and CAP-bound state was inferred from the cryo-EM structures and mutagenesis experiments^20,21^: upon binding, capsaicin establishes a hydrogen bond with the S4-S5 linker thereby “pulling” on this structural element in the outward direction; as in voltage-gated ion channels this motion relieves the constriction exerted by the linker on the S6 bundle and thus triggers dilation at the level of the lower gate.

Despite the fact that the fundamental aspects of capsaicin-triggered activation have been clarified, much remains unexplained concerning the factors affecting the potency of this ligand. Indeed, the ability of TRPV1 to respond to multiple physical and chemical stimuli makes experimental characterization of each gating modality of TRP channels extremely difficult. The challenge comes mostly from the fact that the modalities are allosterically coupled to each other^22–26^: simultaneous application of two or more stimuli in many cases results in cross sensitization. Moreover, it is extremely difficult, if at all possible, to experimentally decompose the process of activation into stimulus sensing, coupling of the sensor to the gate and opening of the gate; thus the phenotype of a single residue mutant is, in general, extremely difficult to interpret. As a result, several seemingly contradictory conclusions have been drawn from experiments concerning, for instance, temperature sensitivity^27–30^ or modulation by lipids^20,31–35^.

To tame this complexity and reveal the molecular mechanism underlying TRPV1 activation a quantitative molecular model of the activation process is desirable whose predictions can be objectively compared with the available experimental data and be used to design new experiments. To this end, we use here state-of-the-art molecular dynamics and free energy calculations. The resulting model is in agreement with the experiments performed so far. Moreover, by providing alternative interpretations of previously published data, we reconcile seemingly contradictory observations about water accessibility of amino acids in the pore domain. Importantly, we generated novel predictions about two single residue mutations, M572A and F580Y, that we confirmed experimentally using electrophysiology.

Anticipating our results, we find that TRPV1 activation relies on the rotation of an evolutionarily conserved amino acid located in the middle of S6 (N676), which project its side chain toward either the central pore or a peripheral cavity. Hydration of the pore and thus conduction of ions is possible only in the former case. We find that the conformational switch of N676 is triggered by a markedly temperature dependent wet/dry transition of the peripheral cavity that favors the conductive state of the channel at high temperature.

## RESULTS

### Four peripheral cavities in the capsaicin-bound state

To examine the differences between the three TRPV1 cryo-EM structures (apo, RTX-bound and CAP-bound), we aligned their S1-S4 domains, which do not undergo significant conformational rearrangements during gating, and compared the superimposed pore domains. Surprisingly, even though the CAP-bound is a putative intermediate state, we found it to be distinct from the other two, *i.e.* it differs from the apo and RTX-bound states more than the latter two differ from each other. Unlike the apo and RTX-bound conformations, in the CAP-bound one the S6 C-terminus part moves away from the S4-S5 linker, suggesting that these two elements are not mechanically coupled as they usually are in channels of the 6TM family (Figure S1A). As noted by Cao et al.^19^, uncoupling is accompanied by the breaking of a hydrogen bond between S4-S5 and S6.

Partial detaching of the S6 C-terminus part from the S4-S5 linker can result in formation of small cavities between these two elements. In fact, using a well established method to detect voids and pockets in proteins (f-pocket^36^), we found four cavities (hereafter referred to as peripheral cavities or PCs) in the CAP-bound state (one per channel subunit) that are connected to the intracellular compartment and run parallel to the central pore up to the level of N676 (the middle of S6) (Figure S1B). Notably, in the apo and RTX-bound states, these cavities are significantly smaller and not connected to the intracellular solution.

### Alternate hydration of central pore and peripheral cavities

The presence of four PCs, accessible to the intracellular solution, in the CAP-bound state of TRPV1 made us hypothesize that these are normally filled with water. Therefore, in modeling the initial configuration for molecular dynamics (MD) simulations, we filled these cavities with several water molecules (from 4 to 6) and considered a system with empty PCs as a control case. For each of them, a trajectory of ∼ 0.75 μs was generated (see Methods). Note that during these runs the PCs remained either hydrated if they were initially filled with water or dehydrated if they were initialized as empty pockets, suggesting that the channel admits two metastable states.

To investigate how the presence or absence of water in the PCs affected the stability of the channel, we analyzed the conformational rearrangements of the pore domain along the MD trajectories using as reference the three cryo-EM structures (apo, RTX-bound and CAP-bound). In particular, we pooled together the configurations sampled during the two 0.75 μs long simulations and the three experimental structures; we then calculated the whole set of pairwise structural distances (root mean square deviation, RMSD, of the position of heavy atoms) to characterize possible structural drifts toward one or another of the cryo-EM structures. To best analyze these distance relations, we represented the configurations as points in the two-dimensional space generated by multi-dimensional scaling (MDS)^37^. As an outcome, the MDS algorithm assigns to each configuration two coordinates with the property that Euclidean distances in this embedding space are maximally similar to the original ones (RMSD). Figure 1 shows that the channel with hydrated PCs fluctuates in the region surrounding its initial structure, *i.e.* the CAP-bound state. This result indicates that the initial structure corresponds to a stable conformational state of TRPV1 and argues against the hypothesis of a conformational average between the closed and open states. Interestingly, the other system, in which the PCs were left empty, drifts away from the CAP-bound state. In this case, two out of four subunits converge to the open state. Similar results were obtained by considering alternative definitions of structural distance (Figure S2).

**Figure 1:**
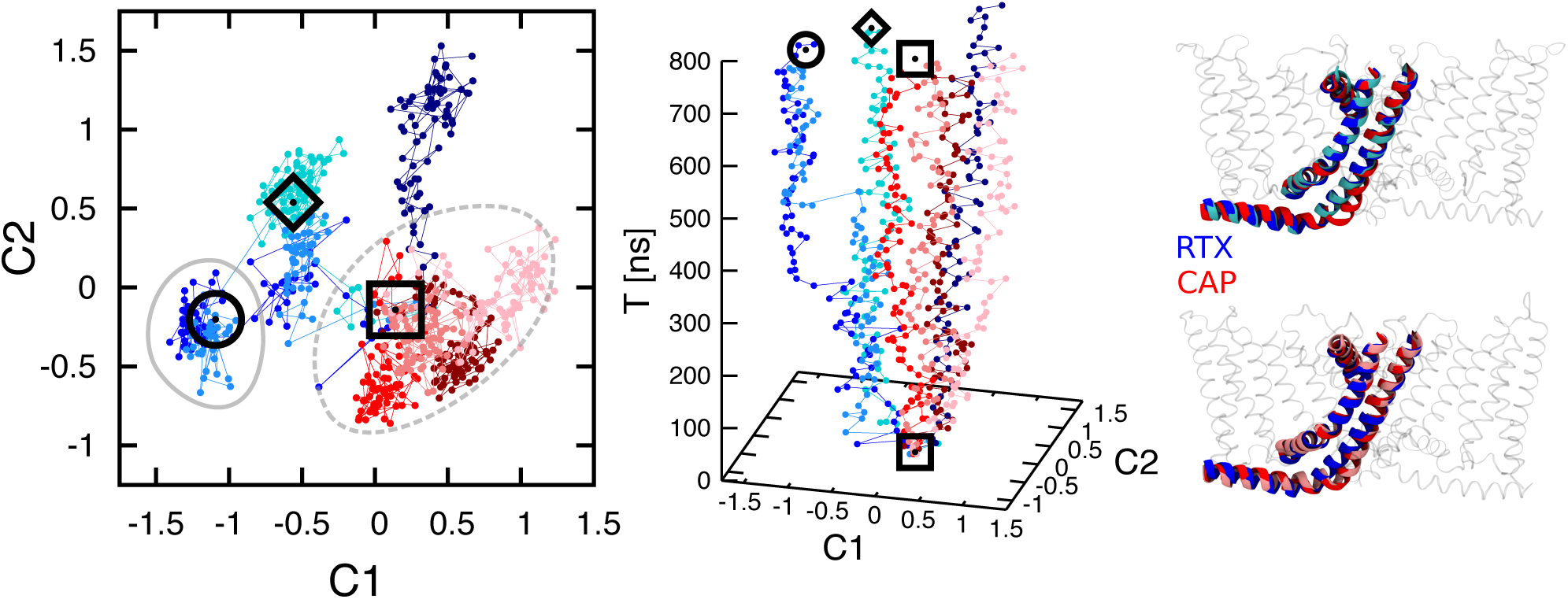
Multidimensional scaling (MDS) analysis of the two MD trajectories of TRPV1 with empty and hydrated PCs. Each configuration is rendered as a point in a plane parameterized by coordinates C1 and C2 (left panel). In the middle panel the time axis is added for clarity. The diamond, square and circle denote the cryo-EM apo, CAP-bound and RTX-bound structures respectively. Each of the colored lines corresponds to one subunit of the channel: blue-shaded and red-shaded lines correspond to the simulations with empty and hydrated PCs respectively. Note that all four subunits with hydrated PCs fluctuate around the initial state (CAP-bound), while two of the subunits with empty PCs converge to the open structure. In the right panel, representative subunits of the two MD trajectories are shown: the channel with empty PCs on the top (the subunit in cyan almost coincides with the RTX-bound structure shown in blue) and the channel with hydrated PCs on the bottom (the subunit in pink almost coincides with the CAP-bound structure shown in red).

Encouraged by these observations, we asked whether the final configurations reached by the simulations with empty and hydrated PCs were, in fact, conductive and non-conductive states, respectively. To answer this question, we characterized the average radius and hydration profiles of the central pore for the two MD runs. Figure 2A shows that the estimated radius profiles are qualitatively similar: the most prominent differences are observed in the central cavity and at the lower gate (in the channel with hydrated PCs the radius of the central cavity is larger and the constriction region of the lower gate is longer). In contrast, the estimated hydration profiles are drastically different: in the channel with hydrated PCs, there are two gaps in the water density profile at the levels of the upper and lower gates that indicate that this conformational state is non-conductive; however, in the channel with empty PCs, the central pore is continuously hydrated, suggesting that both gates are open. Monitoring the time course of hydration (Figure 2B) we found that the opening of the upper and lower gates takes place between 410 and 430 ns. Interestingly, these compartments become hydrated approximately at the same time (after ∼ 410 ns and 430 ns for the lower and upper gates, respectively), a near simultaneity that suggests thermodynamic coupling.

**Figure 2:**
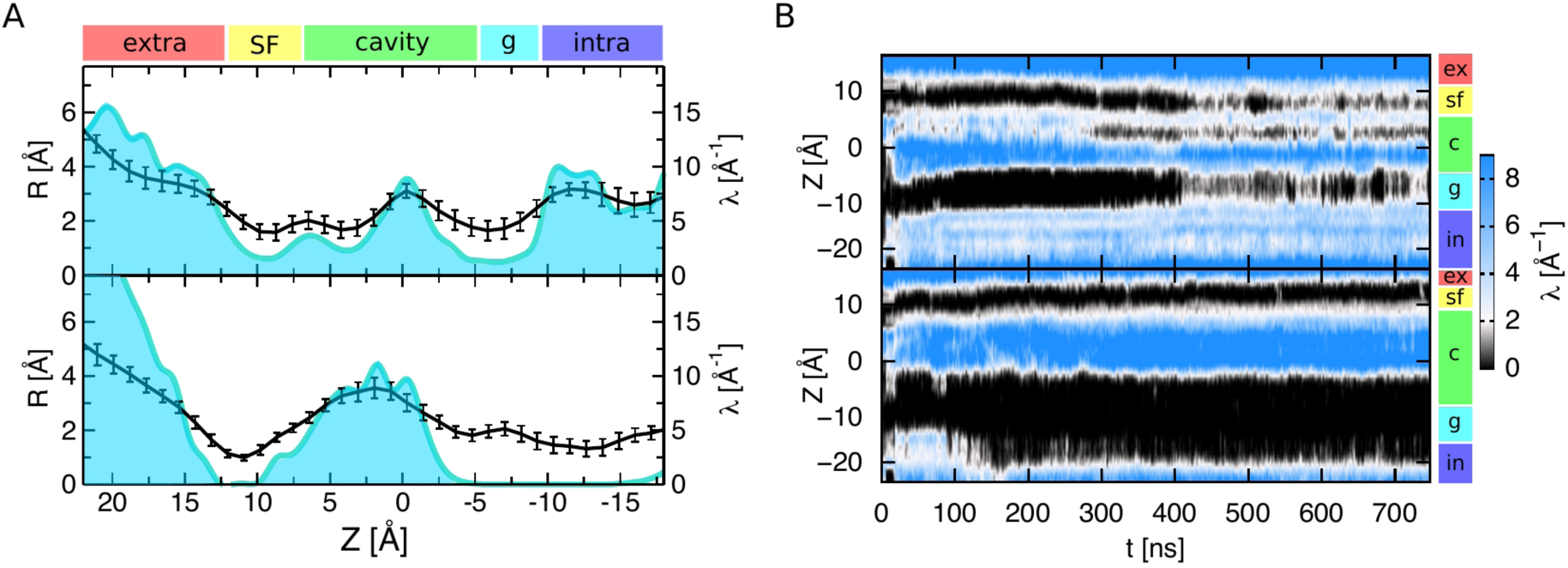
Pore radius and water density profiles for the channel conformations with empty (top panels) and hydrated (bottom panels) PCs. A. The plots show the average radius (black curve) and water density (cyan curve) profiles calculated for the second halves of the MD trajectories. Note that in the case of hydrated PCs the water density profile is interrupted at the levels of the upper and lower gates while it is continuous in the other case. Error bars show the standard deviation. Extra, SF, cavity, g and intra denote the extracellular solution, selectivity filter, central cavity, lower gate and intracellular solution, respectively. B. Time course of water density in the central pore. In the simulation with empty PCs, both the upper and lower gates become hydrated between 410 and 430 ns. In the other conformation, the two compartments remain dehydrated during the entire MD run.

Finally, we estimated free energy profiles for permeation of a sodium ion in the two channel conformations using metadynamics^38^ (Figure 3 and S3; see Methods). We found that, in the channel with hydrated PCs, there is a large free energy barrier at the level of the lower gate, greater than 12 kcal/mol. This indicates that ion permeation is strongly disfavored and thus that this conformation is non-conductive. In the other case, the free energy barrier is of ∼ 5.3 kcal/mol (in the direction from intra-to extra-cellular solutions), indicative of a partially conductive state. The free energy profile for the channel with empty PCs is in qualitative agreement with what previously found for the RTX-bound structure^39,40^. We note that, in the channel with empty PCs, the lower gate barrier is ∼ 2 kcal/mol higher^39,40^. We attribute this difference to the fact that this conformation is not symmetric: only two subunits out of four correspond to the RTX-bound structure (Figure 1). We hypothesize, however, that on longer timescales, the channel with empty PCs would evolve toward a more symmetric conformation, coincident with the RTX-bound structure.

**Figure 3:**
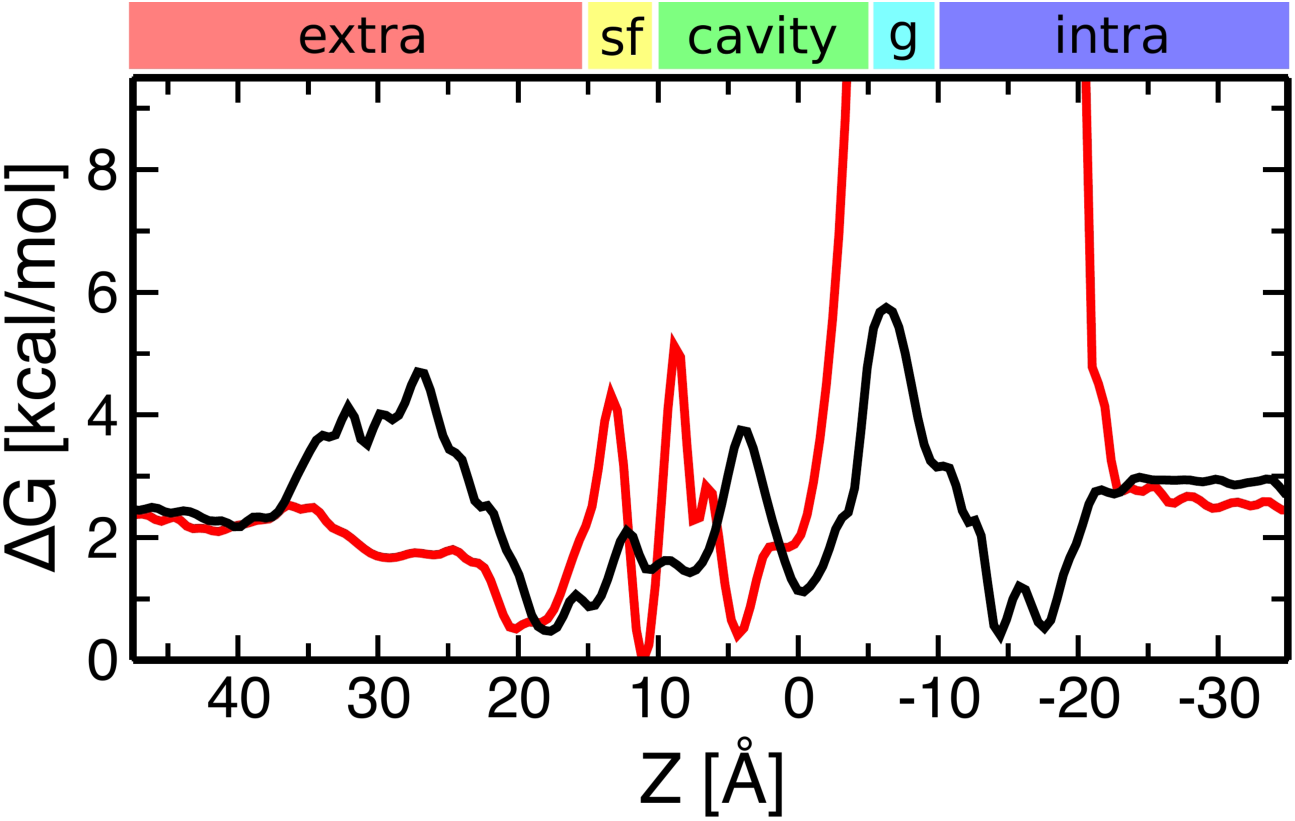
Free energy profiles for permeation of a sodium ion in the conformation with empty (black) and hydrated (red) PCs. Note the large free energy values at the level of the lower gate in the conformation with hydrated PCs indicating that this conformation is non-conductive.

Based on these analyses, we conclude that the conformation with hydrated PCs is closed, while the conformation with empty PCs is partially open. For the sake of clarity, hereafter we will refer to these conformations as closed and open, respectively (however, we will keep them distinct from the cryo-EM closed and open states by using the nomenclature apo and RTX-bound).

### The presence of peripheral cavities reconciles seemingly conflicting experimental observations

In a milestone paper, Salazar et al.^41^ reported an in-depth investigation of TRPV1 pore accessibility to small solutes. These investigators mutated one by one all the S6 residues to cysteine and probed their accessibility to the intracellular solution in the conductive and non-conductive states using covalent modification by several reagents. In the TRPV1 conductive state, all of the S6 residues could be accessed from the intracellular solution. In the non-conductive state, instead, the residues up to the level of I672 and Y671 could be modified by small reagents, and only up to the level of I679 by bulky reagents. Based on these data, the authors suggested that in the non-conductive state the lower gate (I679) is partially dilated, while the upper gate (I672-Y671) remains tightly constricted. This elegant explanation, however, turned out to be inconsistent with the structural information subsequently obtained via cryo-EM on the apo conformation^18,19^. Indeed in this structure, which corresponds to a non-conductive state of the channel, the lower gate is tightly constricted with a radius ∼ 1.0 Å. Moreover, Salazar et al. found that I672 was the last residue accessible from the intracellular solution in the non-conductive state, while in the apo structure the upper gate is much more displaced toward the extracellular compartment, lying between residues G643 and M644.

These seemingly contradictory experimental observations could be reconciled by noting that S6 residues are accessible to water not only through the pore but also through the PCs. Or, in other words, can the puzzling accessibility profiles inferred by Salazar et al. provide experimental evidence toward the existence of PCs? To answer this question, we characterized the accessibility of the S6 residues in the two states of interest, open and closed. As a control case, we considered also the apo structure; for this structure, we performed a long MD run to fully equilibrate the protein in the membrane/solution environment (see Methods). We used, as a measure of water accessibility, the overlap between the atomic density of a given S6 residue and that of water molecules coming from the intracellular compartment (Figure 4; see Methods). In agreement with Salazar et al., we found that in the open state all the residues are accessible to the intracellular solution. Interestingly we found that in the closed state several residues located above the lower gate (I679), in particular L672, L673, L675, N676, M677 and L678, are also accessible; we note, however, that the water molecules interacting with these residues are not located in the central pore but in the PCs. In the apo state (bottom panel), some of these residues (L674 and M677) are also accessible through the PCs. Hence, combining the results obtained for the two non-conductive states (closed and apo), we found that all of the S6 residues between the lower gate and L672 are accessible to the intracellular solution, a result in complete agreement with the observations by Salazar et al.

**Figure 4:**
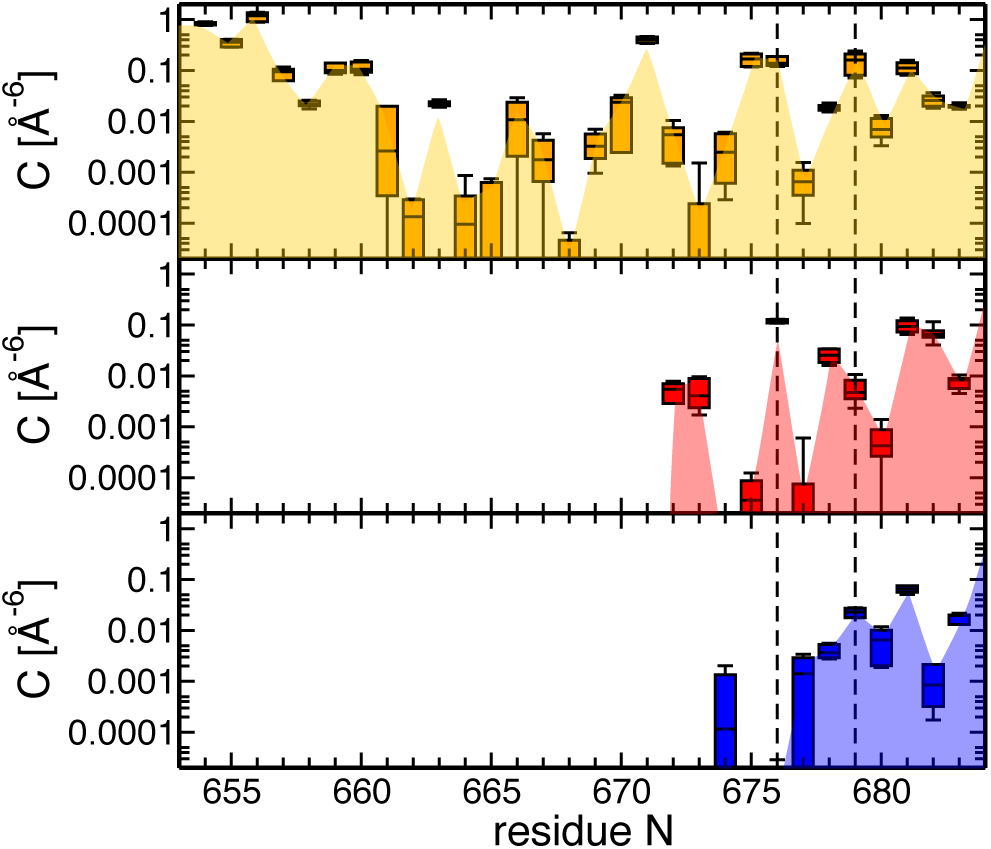
Water accessibility of S6 in the three TRPV1 states, open (top), closed (middle) and apo (bottom). Water accessibility is measured as overlap between the atomic density of a given S6 residue and that of the intracellular water (see Methods; the values shown correspond to a sum over all four channel subunits). In the open state, all S6 residues considered in the analysis are accessible to the intracellular solution; in the closed state, residues up to L672 (except L674) are accessible; in the apo structure, residues up to M677 and also L674 are accessible to the intracellular solution. Error bars show the standard deviation. Extra, SF, cavity, gate and in denote the extracellular solution, selectivity filter, cavity, lower gate and intracellular solution, respectively.

### Evidence for hydrophobic gating

The PCs are located between the S4-S5 linker and the S6 C-terminus of one subunit. The residues lining the cavities are mostly hydrophobic (Figure S4); there are few hydrophilic residues at the entrance of the PCs and only one amino acid deep inside the PCs, namely N676. We noticed that this residue adopts different conformations in the closed and open states (Figure 5). In particular, when the channel is closed, N676 is exposed to the PCs, where it forms multiple hydrogen bonds with the resident water molecules. Interestingly, the central pore is devoid of waters in the region proximal to this residue. In the open state, instead, N676 faces the central pore, which is hydrated throughout its length. In this case, the PCs are occupied by the side chains of M682, which adopt an extended conformation.

**Figure 5:**
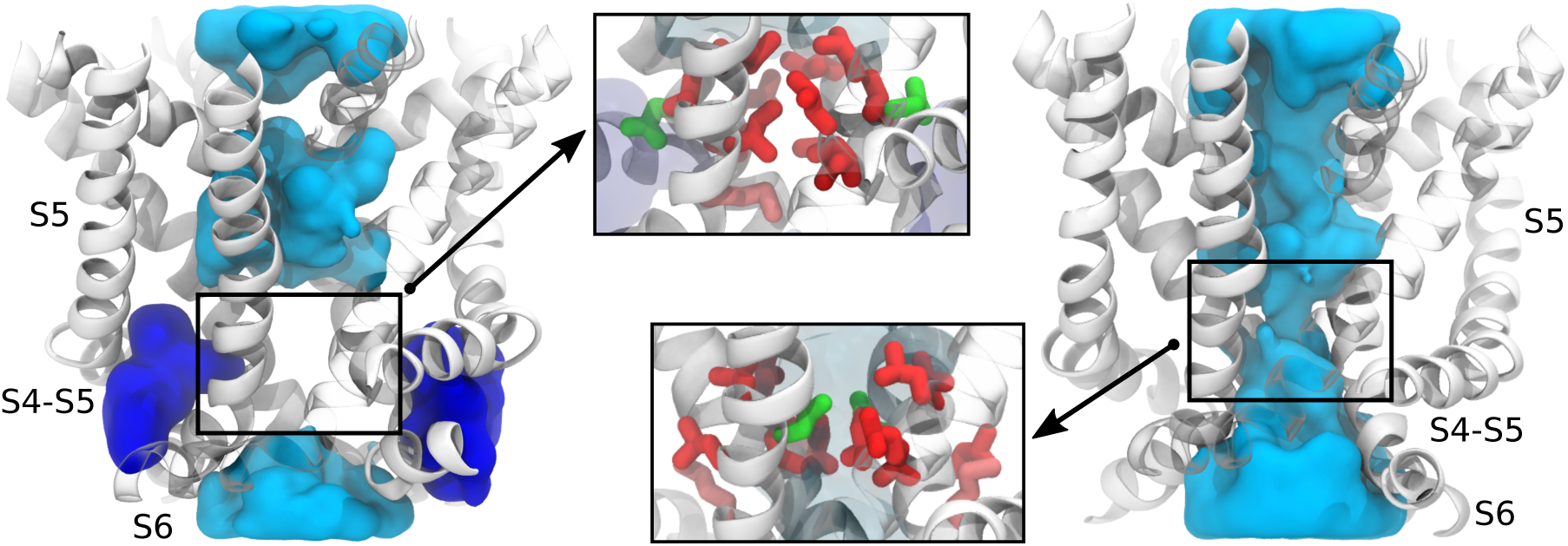
Two different states of the TRPV1 pore domain, hydrated (left) and empty (right) PCs. The PCs are located between the S4-S5 linker and S6. Water in the PCs and in the central pore is shown in blue and cyan, respectively. Note that, in the state with hydrated PCs, there are two interruptions of water density in the central pore: at the level of the upper and lower gates. In the state with empty PCs, the water density is continuous. The two conformations of N676 (green) are shown in the insets: this residue is either facing the PCs (when these are filled with water) or the central pore (when the PCs are empty). The residues of S6 surrounding N676 are shown in sticks and colored according to their polarity^42^ (red – hydrophobic, green – hydrophilic). Note that all of these residues (except N676) are hydrophobic.

As discussed above, the pore radius profiles of the closed and open pores are qualitatively similar (Figure 2A); thus the different hydration in the two cases does not reflect a constriction or dilation of the channel pore, but rather results from subtle changes in the hydrophobic character of the pore. To test this hypothesis, we estimated the per-residue solvent accessible surface area (SASA, see Methods) for the S6 residues (A665-I679) in the closed and open states. Note that only water in the central pore was considered for the analysis. Figure 6 shows that the two SASA profiles are significantly different. In particular, the upper part of S6 (T670, Y671, L674 and L675) is more hydrated in the closed pore compared to open, while the N676 and I679 residues of the lower part of S6 have a larger SASA when the channel is open. It is important to note that, in the closed state, N676 has zero SASA; this is consistent with our previous observation that N676 is facing the PC, not the central cavity. These data indicate that, in transitioning from the closed to open pore, the SASA of several hydrophobic residues (L674, L675, L678, and also Y671) decreases while the SASA of hydrophilic N676 increases, suggesting that overall the open pore is more hydrophilic than the closed one.

**Figure 6:**
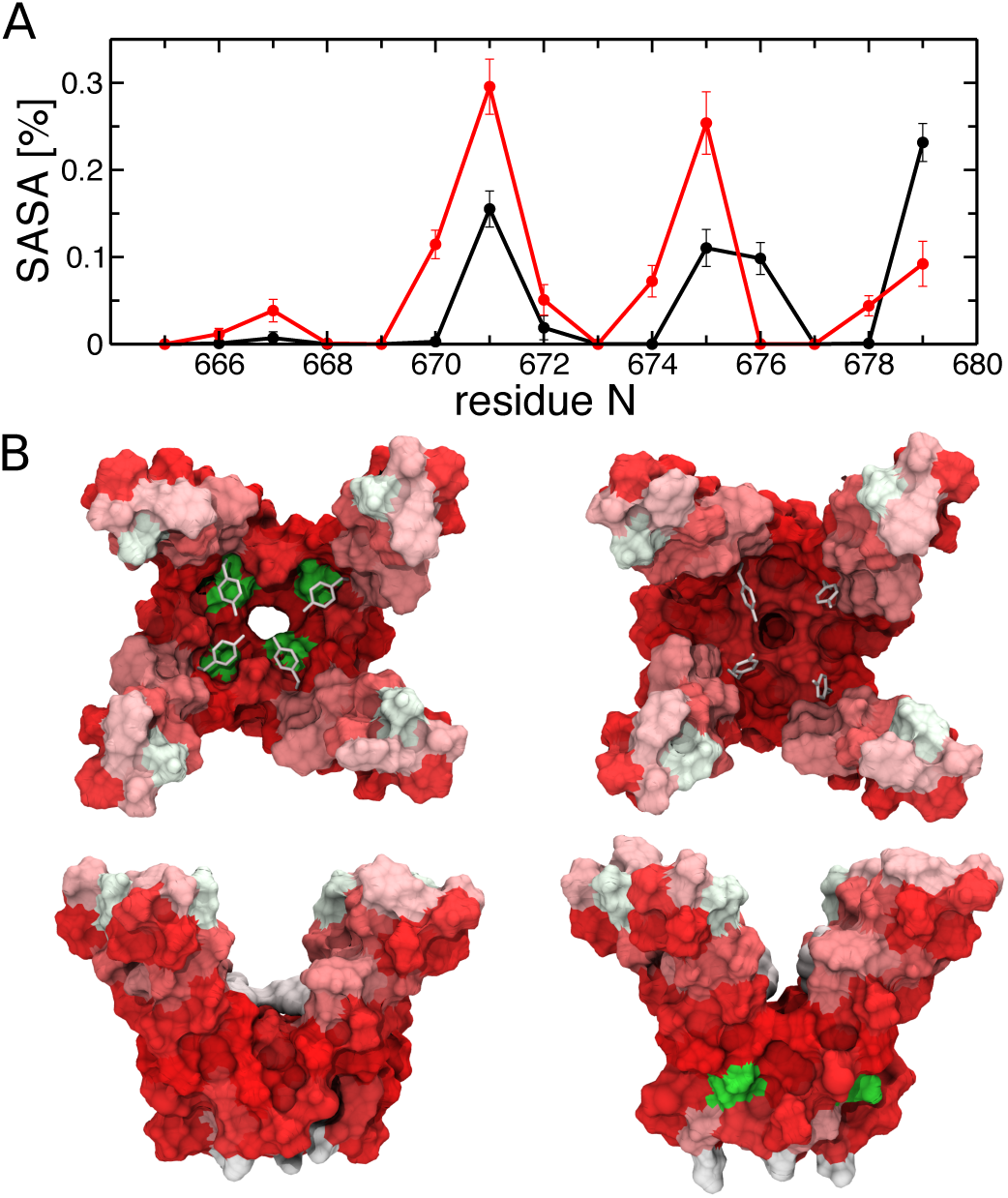
Exposure of the S6 residues to the channel central pore. A. Solvent accessible surface area (SASA) of the S6 residues estimated for the open (black) and closed (red) states. Only water in the central pore was considered for the analysis (see Methods). The N-terminus part of S6, positions 665-675, is more hydrated in the closed state than in the open one, while two C-terminus residues, N676 and I679, have larger SASA in the open than in the closed state. Error bars show standard deviations. B. Polar properties of the inner and outer surfaces of the S6 bundle in the open (left) and closed (right) states: top (top panel) and side (bottom panel) views. The residues are colored according to polarity^42^ (red – hydrophobic, and green – hydrophilic). In the open state, there is a polar (green) spot in the inner surface of the S6 bundle, *i.e.* facing the central pore; this spot corresponds to N676. In the closed state, the inner surface of the S6 bundle is entirely hydrophobic; in this case, N676 projects its polar group toward the outer surface of the bundle, *i.e.* outside the central pore.

### Temperature-dependent mechanism of activation

To assess the stability of the TRPV1 states obtained via MD simulations, we estimated the free energy profile of the closed-open transition for a single subunit using metadynamics^38^. As a collective variable (CV) we used the orientation of N676 with respect to the central pore. In particular, we considered a local reference frame at the crossing point between two S6 helices on the membrane plane (xy) defined by the projection of one of the S6 helices’ axis (x’) and its orthogonal direction (y’); we then calculated, for the adjacent S6, the y’ coordinate of the N676 side chain carboxamide oxygen (Figure S5). Note that this collective variable is different from the previously used one (RMSD of the pore domain), which allows to track conformational changes between the closed and open states. To investigate whether the free energy profile is temperature-dependent, we performed metadynamics simulations at three different temperatures: room temperature (300 K), high temperature (340) and, as a control, low temperature (280 K). Starting from the closed state, we explored thoroughly the configuration space to observe transitions in one of the four subunits during 400 ns of simulation.

Figure 7A shows the three estimated free energy profiles (see also Figure S6 for their time evolution). All of them have a local minimum at ∼ -2 Å, corresponding to a conformation with N676 facing the PC. Importantly, a free energy minimum at positive values of the CV (*i.e.* N676 pointing toward the central pore) is present only at 340 K, indicating that the open state becomes more stable as the temperature is increased. Note that in our case the difference between the closed and open states does not exceed 1 kcal/mol, and the barrier separating these states is only of ∼ 1.5 kcal/mol. This indicates that at 340 K the closed and open states are both significantly populated and in fast exchange.

**Figure 7:**
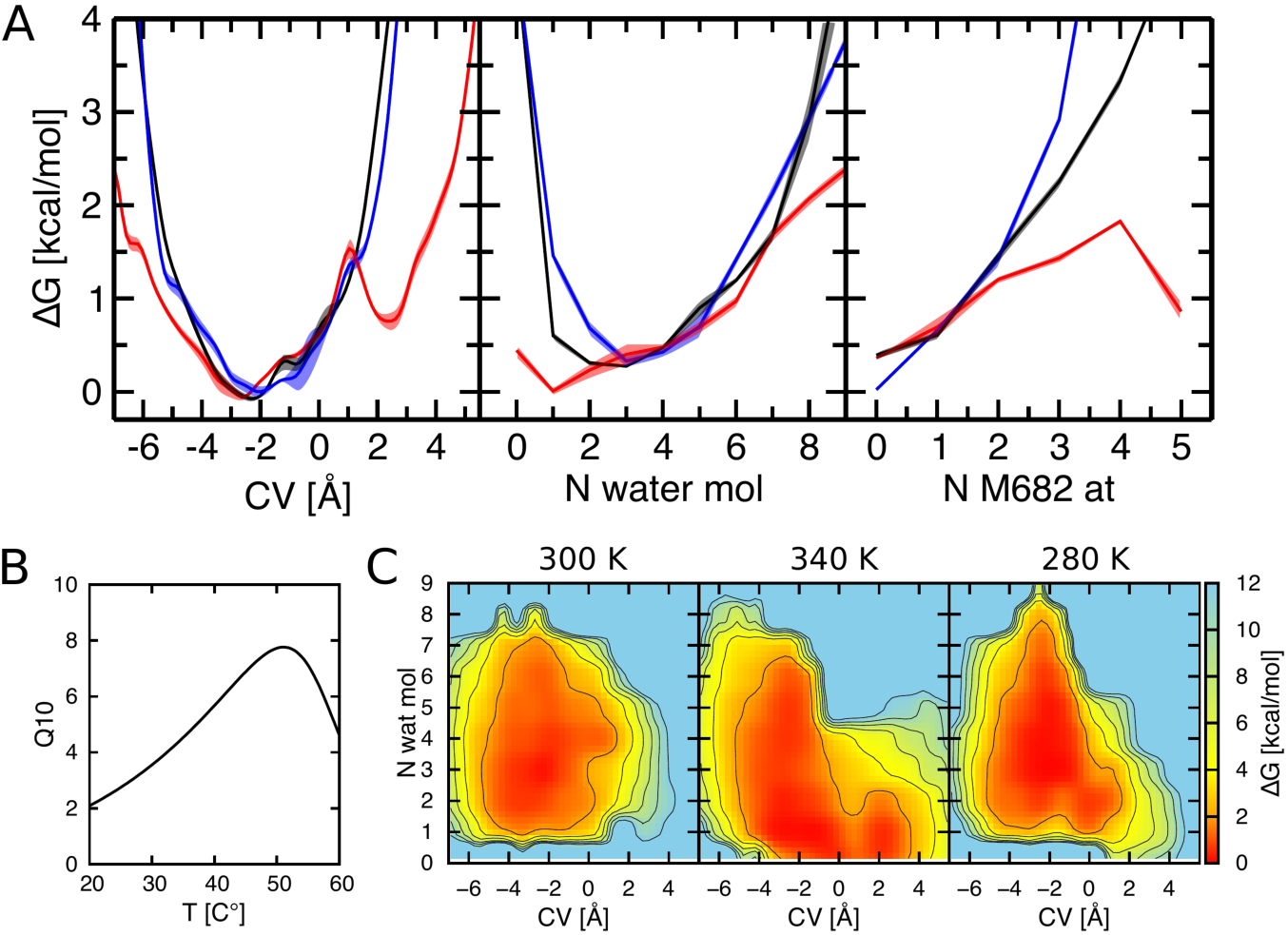
Metadynamics simulations results. A. Free energy profiles (1D) as a function of the N676 orientation (left panel) and of the two non-biased collective variables: the number of water molecules/M682 atoms inside the PC (middle and right panels, respectively). The black, red and blue lines correspond to the average free energy profiles at 300 K, 340 K and 280 K, respectively. The standard errors (see Methods for details) are shown as a shade around each profile. Note that the 300 K and 280 K free energy profiles are very similar to each other. B. Temperature coefficient Q_10_ vs. temperature estimated using the free energy profile of the PC hydration. C. Free energy surfaces as a function of N676 orientation (x-axis) and number of water molecules inside the PC (y-axis). The left, middle and right panels correspond to the simulations at 300 K, 340 K and 280 K, respectively.

Despite the fact that only one out of four N676 residues was subject to the biasing potential, the orientation of this sidechain toward the central pore promoted the re-orientation of remaining ones (Figure S7); indeed rotation of a single sidechain promotes hydration of the central pore, which, in turn, favors the rotation in the remaining subunits. Experimental evidence of this cooperative effect has been previously reported and suggested to be the determinant of temperature sensitivity^43,44^. This thermodynamic coupling among different subunits is likely to stabilize the open state at room temperature, as we indeed observe in our MD simulations of the channel conformation with all four N676 residues facing the central pore.

Besides the biased CV, we estimated the free energy profiles for the hydration of the PC and its occupation by M682 using the reweighting scheme by Tiwary and Parrinello^45^ (Figure 7A). Interestingly, these collective variables are sensitive to high temperature as well. In particular, at 340 K the channel states with the hydrated and dehydrated PC coexist, while at 300 and 280 K the hydrated state is predominant. Additionally, at 340 K, M682 can be found inside or outside the PC with relatively equal probability, while at 280 and 300 K the “outside” conformation is strongly favored. Overall, at 300 K and 280 K the PC tends to be filled with water, whereas at 340 K the PC can be equally occupied by either water or M682.

We further calculated the temperature coefficient Q_10_ in order to make a direct connection to experiments. To estimate the free energy difference between the open and closed states at the three different temperatures, we used the free energy profile of the PC hydration (see SI for a detailed explanation). The calculated Q_10_ (Figure 7B) is in quantitative agreement with what previously found for the TRPV1 turret deletion mutant^26^ (*i.e.* the same construct considered here). We conclude that the suggested activation mechanism shows temperature-dependent features that are intriguingly compatible with experimentally observed temperature sensitivity of TRPV1. We note, however, that the mechanism of TRPV1 temperature sensitivity is possibly more complex and involves also other channel regions such as the N-terminus^27,46,47^, the C-terminus^28,48,49^, the extracellular loops^30,50^, the transmembrane domain^51^ and the TRP box^52,53^.

Finally, we estimated two-dimensional (2D) free energy surfaces for pairs of CVs such as the orientation of N676 with respect to the central pore and the PC hydration (see Figure S8A for other pairs). Figure 7C shows that at 300 K and 280 K the channel is mostly found in one state, in which N676 faces the PC and the latter is filled with water. At 340 K, however, this state coexists with the other one, in which N676 faces the central cavity, and the PC is either dry or filled with one water molecule only. These data indicate that at 340 K the hydration/dehydration of the PC and the orientation of N676 are correlated. Therefore hydration of the PCs controls the orientation of N676 and thus, ultimately, pore conductance.

### Mutation of residues facing the peripheral cavities affects TRPV1 activation

To test our molecular models we designed TRPV1 mutants with altered thermodynamic equilibrium between the closed and open states. In particular, we mutated M572 and F580, facing the PC, into alanine and tyrosine, respectively. We measured the responses of the obtained mutants to capsaicin, low pH and heat using two-electrode voltage clamp (TEVC) and Ca^2+^ imaging experiments and compared them to those of the wild type (WT) channels. According to the model suggested here, mutation of M572 and F580 should stabilize the closed conformation by favoring the PC hydration. Importantly, both of them are located far from the vanilloid binding site and the residues involved in pH sensing^54–58^, and therefore a large effect on the capsaicin-or low pH-evoked responses should indicate an impaired coupling between the protein elements affected by these stimuli and the activation gate. Moreover, in the case of F580Y, the structural perturbation entailed by the mutation is minimal, since it only consists in the addition of a hydroxyl moiety.

Figure 8A-D shows that the capsaicin dose-response curve of the F580Y mutant is shifted toward higher concentrations of capsaicin (Figure 8C and D). In the case of M572A, however, this curve was similar to that of the WT (Figure 8B and D). Both mutants showed reduced sensitivity to low pH compared to the WT. The F580Y mutant specifically responded only to pH 4 but not to lower proton concentrations (Figure 8E-H). Finally, we measured heat-induced Ca^2+^ responses of the M572A and F580Y mutants and compared them to that of the WT (Figure 8I-M): in practice, we stimulated the channels with a heat pulse, followed by the application a low pH solution (pH = 4) and 0.5 μM capsaicin. Figure 8I, J and M shows that the M572A mutant had reduced heat sensitivity, while the activation by either low pH or capsaicin was similar to that of the WT. In the case of F580Y, the Ca^2+^ responses evoked by either heat or capsaicin were significantly smaller than those of the WT, while the activation by low pH was not affected (Figure 8I, K and M). Control HEK cells transfected with an empty vector showed very small transient Ca^2+^ responses at pH 4 and no response to either heat or capsaicin (Figure 8L). Note that for these measurements we used a low affinity variant of Fura-2 to avoid saturation of the dye by large Ca^2+^ currents due to TRPV1 activation. Overall, reduced responses of M672A and F580Y to the two and three tested stimuli respectively indicate that the introduced mutations resulted in stabilization of the closed conformation of TRPV1, confirming our predictions made based on the molecular modeling. We also generated three additional mutants, F580A, N676T and N676V, which should show altered activation according to our model. These mutants showed no clear responses to either low pH (pH=4) solution, or capsaicin concentrations as high as 300 μM in TEVC experiments, thus we could not characterize them.

**Figure 8:**
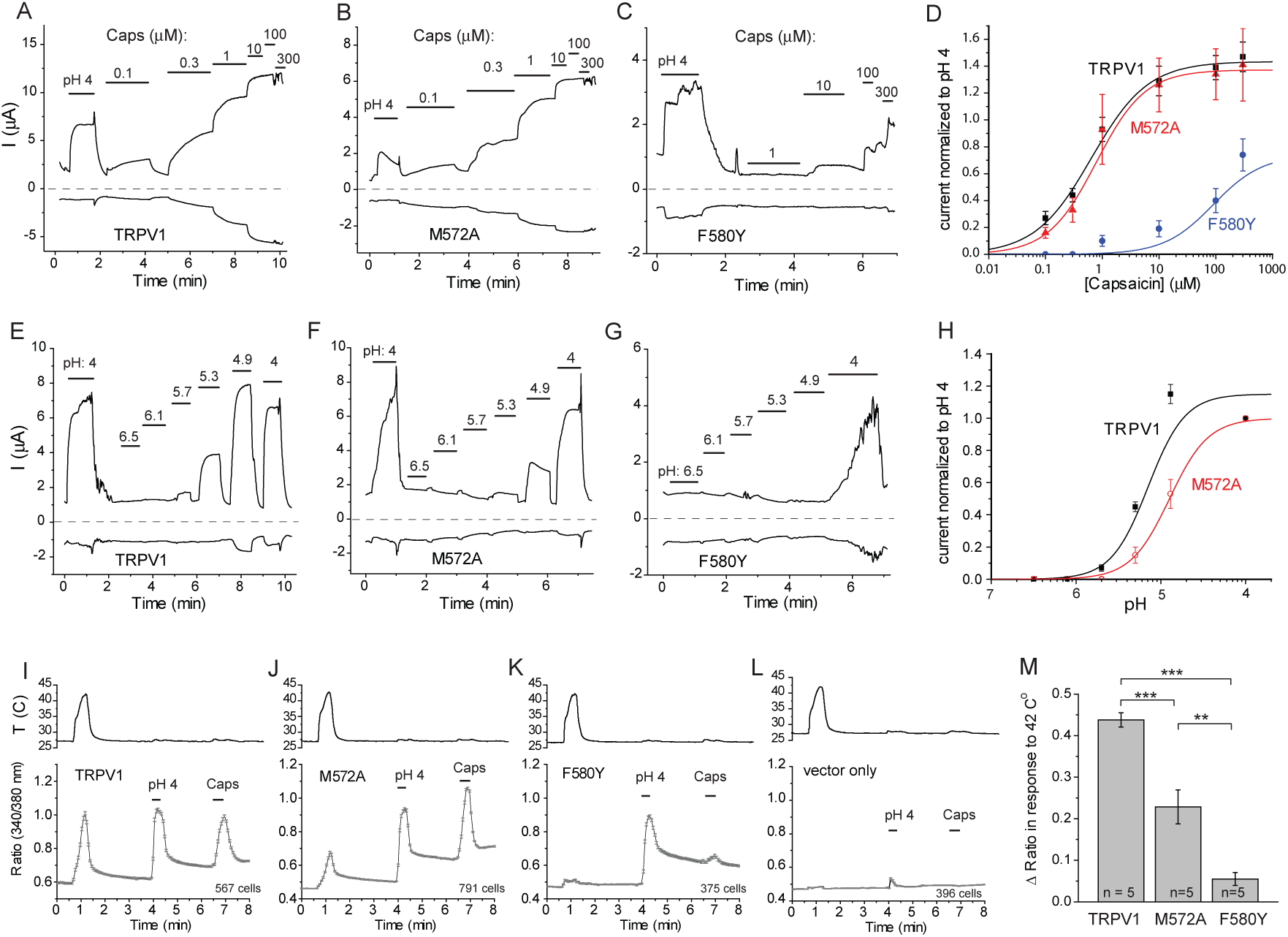
Experimental characterization of the M572A and F580Y mutants of TRPV1. A-C. Representative current traces in response to different capsaicin concentrations of the WT (A), M572A (B) and F580Y (C). TEVC oocyte electrophysiology was performed as described in Methods. Currents at 100 mV and -100 mV are shown. Zero current is indicated by the dashed-line. Oocytes expressing the WT or the mutants were stimulated first with a pH 4 solution and then with successive concentrations of capsaicin. D. Hill fits of the WT and the two mutants responses to different capsaicin concentrations at 100 mV. Currents were normalized to that evoked by pH 4. E-G. Representative current traces in response to low extracellular pH of the WT (E), M572A (F) and F580Y (G). H. Hill fits of the WT and the M572A mutant responses to solutions with different pH solutions. Currents were normalized to that evoked by pH 4. I-L. Intracellular Ca^2+^ responses to heat, pH 4 and 0.5 μM capsaicin of the WT (I), M572A (J), F580Y (K) and HEK cells transfected with an empty vector (L). Intracellular Ca^2+^ concentration was measured with low affinity Fura-2 as described in Methods. The traces show average +/- SEM of cells from 5 coverslips (the number of cells is indicated in each panel). Top panel shows average temperatures in the vicinity of cells measured during the experiments using a temperature probe and the pClamp software. M. Statistical summary of the response to heat. The average responses of cells from each coverslip were treated as individual data points; n corresponds to the number of coverslips. Statistical significance was calculated with one-way ANOVA, and Bonferoni post hoc comparison.

## DISCUSSION

The structure of TRPV1 in complex with capsaicin determined via cryo-EM is in a non-conductive state. Indeed, the selectivity filter appears to be constricted compared to the structure in complex with resiniferatoxin and DkTx^19^. Moreover, we show here that the lower gate (at the level of I679) is dehydrated and hence impermeable to ions. These observations suggest that TRPV1 is closed at cryogenic temperatures even at high capsaicin concentrations. Consistently, Jara-Oseguera et al.^26^ have shown that this particular TRPV1 construct (turret deletion mutant) populates equally the open and closed states at room temperature (in presence of capsaicin) and becomes increasingly more closed as the temperature is decreased. Taken together, these observations indicate that the CAP-bound conformational state (as determined by cryo-EM) should undergo a temperature-dependent transition toward a conductive conformation that is possibly similar to the open (RTX-bound) state. Our study shed light on this transition by investigating the pathway and the corresponding free energy profiles for different values of temperature.

We found that the lower gate is open or closed depending on the conformation of N676, an amino acid located on S6 at the edge of a π-helix segment. This segment is characterized by extreme conformational flexibility^59^: not all the backbone hydrogen bonds can be simultaneously satisfied and therefore their pattern is dynamic. The presence of the π-helix allows N676 to easily rotate in and out the central pore. This motion is, in turn, controlled by the hydration state of the adjacent PC. Besides extensive accessibility experiments by Salazar et al.^41^, the presence of these cavities is supported by alanine scanning mutagenesis performed on the S6 segment^60^; the residues lining the PCs were shown to produce the greatest perturbation to the channel activation in response to several stimuli, including capsaicin and heat.

The peculiar *i-i+5* hydrogen bond pattern results in an “unsaturated” backbone carbonyl group (π-bulge) that does not exhibit a hydrogen bond with any backbone amino group. Importantly, the N676 rotation causes a relocation of this π-bulge between the two S6 adjacent residues (Figure 9A). In particular, in the open state, the π-bulge is on I672 and forms a hydrogen bond with the carboxamide moiety of the N676 side chain, which faces the central pore. In the closed state, instead, the N676 side chain points toward the PC and hence cannot establish a hydrogen bond with the π-bulge on I672. Therefore, the π-bulge moves to the neighboring Y671 residue.

**Figure 9:**
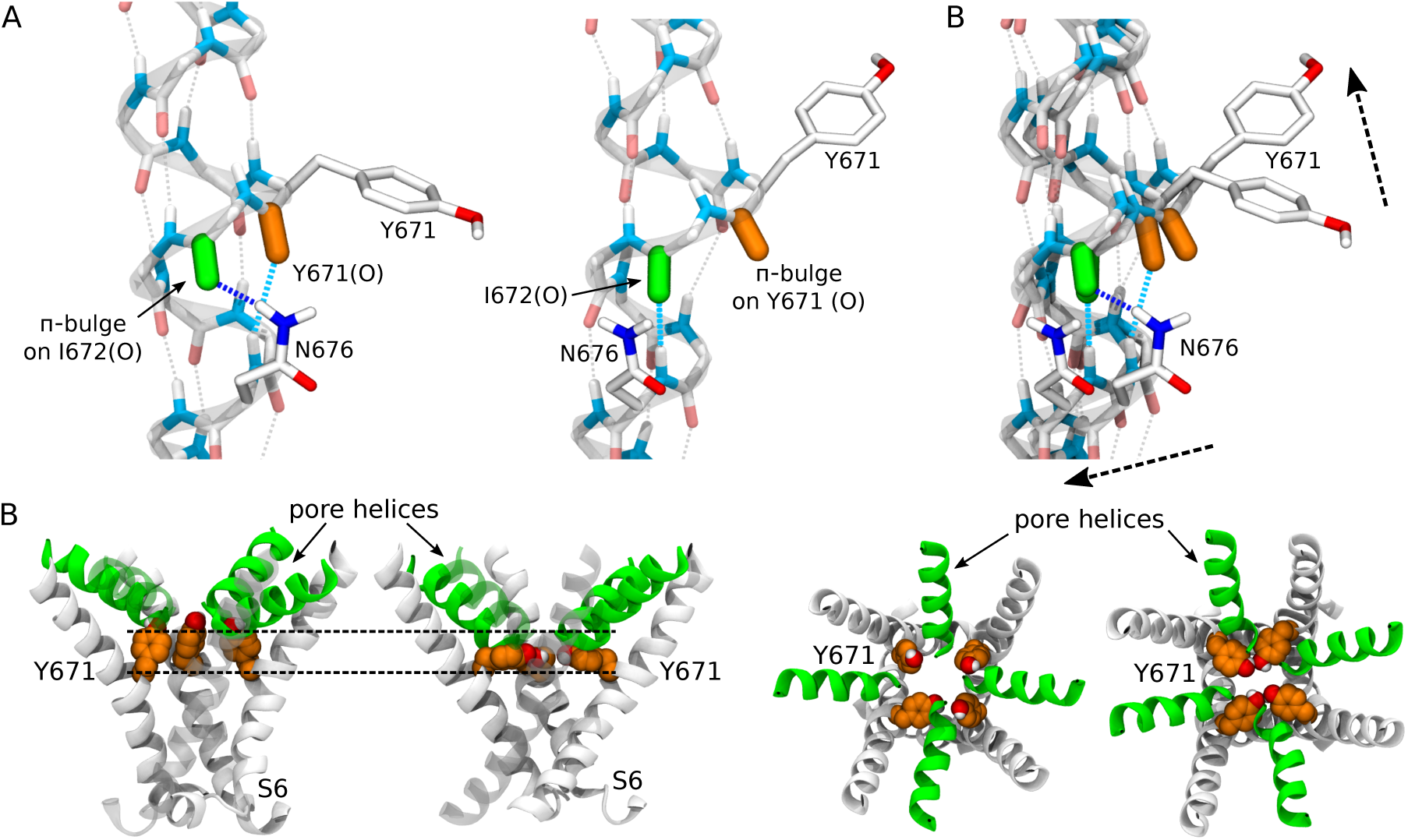
Allosteric coupling between the upper and lower gates of TRPV1. A. Conformational rearrangements involving N676, Y671 and the π-bulge. The left and middle panels correspond to the open and closed states, respectively. The right panel shows the superimposition of the two states. N676 and Y671 are shown as sticks; the carbonyl groups of I672 and Y671 are colored in green and orange, respectively. In the open state, the π-bulge is on I672, which forms a hydrogen bond with the side chain of N676 (shown in blue). The carbonyl group of Y671 is involved in a hydrogen bond with the backbone of N676 (shown in cyan). In the closed state, the π-bulge is on Y671, while I672 forms a hydrogen bond with the backbone of N676. In the right panel, the arrows show the direction of the Y671 and N676 conformational rearrangements during the open to closed transition. B and C. Y671 rotameric states in the closed (left) and open (right) states. B and C correspond to the side and top views, respectively. The Y671 residues, the pore helices and S6 are shown in orange, green and white, respectively. The dashed lines indicate the level of the Cα atoms of Y671 and the lower point of the pore helices in the closed state.

The π-bulge displacement between I672 and Y671 has crucial consequences on the TRPV1 selectivity filter (Figure 9B). Indeed, due to the absence of a restraining hydrogen bond on the carbonyl oxygen, Y671 rotates in the plane of the central axis, a motion that results in a different rotameric state of the side chain of this residue. Thus, while in the open state the phenyl groups are oriented perpendicular to the pore axis and engaged in mutual hydrogen bonds, in the closed state, the same groups are almost parallel to the pore axis. This conformational transition results in a change in Y671 hydration that is fully consistent with very recent fluorescent measurements on a non-natural amino acid introduced at this position^61^. Importantly, in the conformation parallel to the pore axis, the Y671 side chains displace the pore helices by “pushing” in the upward direction (toward the extracellular solution), causing a constriction of the selectivity filter, which disfavors ionic and water transport (Figure 2, 3 and 9B). Overall, the effect of the N676 rotation on the Y671 conformation rationalizes the allosteric coupling between the two gates, a crucial aspect of this channel, in which distinct stimuli can act synergistically^23^.

Interestingly, our molecular mechanism does not entail any large conformational rearrangement of the TRPV1 central pore, whose radius profile is not dramatically altered by the closed-open transition. The hydration/dehydration of this compartment is, in fact, controlled by N676, which upon rotation changes the hydrophobic character of the molecular surface lining the central pore. This susceptibility to perturbations is not uncommon in the pores of ion channels. Wet to dry transitions have been reported several times^62–70^ and are, arguably, the result of a precise evolutionary optimization. Due to a fine-tuned hydrophobicity, pores undergoing hydrophobic gating are poised to transition and therefore are maximally responsive to activating stimuli. In the case of TRPV1, the functional requirement of a polar side chain with alternate exposure to the central pore resulted in clear statistical signals in the set of sequences encoding for this channel. First, an asparagine is present in virtually all the members of the family in the corresponding position^71^. Second, the presence of a π-helix segment involves a unique set of residue-residue contacts across S5 and S6; these contacts (that are observed in the cryo-EM structures) can be inferred for all the members of the family through the co-evolutionary signals detected in large multiple sequence alignments^71^.

Independent experimental structures of different members of the TRP family already highlight the extreme conformational heterogeneity of N676, whose orientation with respect to the central pore can differ by as much as 180° between two structures (Figure S11A). This observation brings additional support to our gating model and raises the possibility that the same molecular mechanism could be at work in other evolutionarily related channels. Strikingly, a recently determined high-resolution structure of a bacterial voltage-gated sodium channel revealed structural waters in regions coincident with the peripheral cavities discussed here^85^; these waters establish hydrogen bonds with the side chain of the asparagine corresponding to N676 in TRPV1 (Figure S11B). Whether these hydrated cavities also trigger a switch between conductive and non-conductive states in voltage-gated sodium channels (which, we recall, share significant sequence features with the TRP family^86^) remains to be confirmed.

An important aspect of our molecular model is that the N676 conformation is controlled by the hydration of a small non-polar cavity. Hydrophobic cavities of precisely this size (hosting few water molecules) have been extensively investigated in the past in model systems, including carbon nanotubes^72–78^. Hydration of these enclosures shows a clear bistable (wet/dry) behavior with resulting equilibrium properties that are highly sensitive to chemical modifications and changes in environmental factors. In particular, hydration has been shown to be exquisitely temperature-and pressure-dependent^79^ and entails a large change in the heat capacity (in analogy to a liquid-vapor phase transition). This feature suggests the possible physical basis for the temperature dependence of TRPV1 gating that we observe in our simulations. As pointed out already by Clapham and Miller in a pioneering work, such variations in heat capacity constitute the likely mechanism of temperature sensitivity in ion channels^80^. Although we cannot exclude that several other molecular mechanisms are at work during temperature-driven activation, we note that according to our free energy calculations, the open probability shows an almost two-hundredfold increase on passing from 300 K to 340 K.

Importantly, the hydration/dehydration transition in small hydrophobic cavities is critically affected by several factors such as pressure, osmolarity and electric fields. This raises the intriguing hypothesis that a unified mechanism of activation might underlie several seemingly unrelated gating modalities. For instance, TRPV1 has been shown to be involved in detection of osmotic stress^81^. Our model can potentially rationalize this observation: an increased concentration of solutes makes segregation of water molecules into the cavity less favorable, thereby promoting dehydration and, ultimately, activation. Another potential implication of the dehydration-triggered activation is the ability to respond to changes in the intracellular hydrostatic pressure as waters can be more or less “pushed” into the cavity. Interestingly, TRPV1 and TRPV4 have been shown to detect exactly these changes as a part of a feedback mechanism that is at work in surface cells of the ocular lens^82^. Moreover, TRPV1 (and also TRPA1) plays a crucial role as airway chemosensor: it activates as a result of binding of single ring benzene derivatives, a possible indication that the cavity can act as non-specific binding site for small hydrophobic moieties^83,84^. Finally, the proximity of the PCs and the predicted PIP_2_ binding site^31^ raises the interesting question as to whether or not this lipid can locally perturb the properties of water and hence modulate TRPV1 activation by affecting the PCs hydration.

## METHODS

### Modeling the initial configuration for molecular dynamics simulations

The structure of the TRPV1 capsaicin-bound (CAP-bound) state was taken from the protein data bank: the pdb code is 3j5r^19^. The structure was refined, and the missing residues were modeled using Rosetta software^87^. Four capsaicin molecules were docked following the protocol described in^88^. The protein with the ligands was embedded in a hydrated 1-palmitoyl-2-oleoylphosphatidylcholine (POPC) bilayer and surrounded by 150 mM NaCl solution. The overall size of the system was ∼ 170x170x160 Å^3^; the total number of atoms was ∼ 400,000. Two MD trajectories were generated with the peripheral cavities (PCs) either empty or hydrated. In the latter case, from 4 to 6 water molecules were added inside each PC. The CHARMM36 force field^89^ was used to describe the protein and the POPC lipids. For capsaicin, we used the parameters derived in^88^. The TIP3P model was used to describe water^90^. An analogous setup was used to simulate the TRPV1 apo state (pdb code 3j5p^18^).

### Molecular dynamics simulations

The equilibration of the systems (three in total: the CAP-bound state with empty and hydrated PCs, and the apo state) was performed using NAMD 2.10 software^91^ in several steps. During the first 4 ns, the temperature was set to 340 K to melt the POPC tails; at this stage, the protein, ligands, water in the PCs, and the POPC headgroups were restrained to their initial positions (harmonic potential, force constant 1 kcal/mol). Then the temperature was set to 300 K, and the restraints were released first from the POPC headgroups (4 ns) and then from the protein side chains after 4 additional ns. In a subsequent stage, which lasted for 100 ns, the remaining restraints on the protein backbone, ligands and water inside the PC were gradually released (by varying the elastic constant from 1 to 0 kcal/mol/Å^2^). Finally, the fully released systems were equilibrated for ∼ 650 ns in the case of the CAP-bound state (with empty or hydrated PCs) and ∼ 200 ns in the case of the apo state.

Simulations were performed at constant temperature and pressure (1 atm) using the Langevin piston approach. For the vdW interactions, we used a cutoff of 11 Å with a switching function between 8 and 11 Å. The long-range component of electrostatic interactions was calculated using the Particle Mesh Ewald approach^92^ using a cutoff for the short-range component of 11 Å. The equations of motion were integrated using a multiple time-step algorithm, with a time step of 2 fs and long-range interactions calculated every other step.

### Metadynamics simulations

#### Na^+^ permeation through the central pore

Metadynamics simulations were performed using the collective variable module implemented in NAMD 2.10^93^. As a collective variable, we considered the z component of the position vector of a Na^+^ ion. The origin was set at the center of mass of the pore domain (residues 562-599 and 657-701). As initial configurations for the closed and open states, we used the final frames of the corresponding equilibration trajectories. The following parameters were considered for metadynamics: width of the Gaussians – 0.25 Å; deposition rate – every 2 ps; height of the Gaussians – 0.001 kcal/mol. The simulations were run for ∼ 1.2 μs in the open state and ∼ 0.7 μs in the closed state.

#### Closed-open transition of one channel subunit

Metadynamics simulations were performed using Gromacs 5.0.4^94^ and Plumed 2.1.5^95^ at three different temperatures: 300, 280 and 340 K. For each of them, the initial configuration was prepared by extracting the final frame from the closed-state equilibration trajectory and relaxing this frame during 5 ns at the corresponding temperature. As a collective variable (CV), we used the orientation of N676 with respect to the central pore: we considered a local reference frame at the crossing point between two S6 helices on the membrane plane (xy) defined by the projection of one of the S6 helices’ axis (x’) and its orthogonal direction (y’); we then calculated, for the adjacent S6, the y’ coordinate of the N676 side chain carboxamide oxygen (Figure S5). Using this CV, we sampled for 400 ns and at different temperatures the transition between the closed state and a state with three subunits closed and one open. The following parameters were used for the simulations: width of the Gaussians – 0.03 nm; deposition rate – every 4 ps; the height of the Gaussians was varied along the simulation and set to 0.1 kJ/mol for the first 200 ns, 0.05 kJ/mol for the subsequent 100 ns and 0.025 kJ/mol for the remaining 100 ns. In all the three simulations the biased CV sampled the entire configuration space and made several transitions between two major conformations: with N676 facing the central pore or the PC (Figure S15B).

In order to keep temperature (300, 340 or 280 K) and pressure (1 bar) constant during metadynamics simulations we used Nose-Hoover thermostat^96^ and Parrinello-Rahman barostat^97^. For the vdW and electrostatic interactions we used a setup similar to that used for the equilibration.

### Analysis of molecular dynamics and metadynamics trajectories

#### Multidimensional scaling (MDS) analysis of RMSD and distance maps overlap data

To compare the protein conformations extracted from the equilibration trajectories with each other and to the three cryo-EM structures^18,19^, we calculated, for each pair of configurations, root mean square deviation (RMSD) and distance maps overlap. First, from every trajectory we extracted 750 protein conformations (using a stride of 1 ns. We then added to the pool of configurations the apo, CAP-bound and RTX-bound structures and calculated the two distance matrices. For the RMSD, we considered the backbone of the pore domain: residues 562-599 and 657-701. For the distance maps overlap, we used the same set of residues, however, added to the analysis their side chains. The distance maps were calculated using gmx mdmat program of Gromacs 5.0.4 toolkit^94^. Finally, we applied multidimensional scaling (MDS) analysis to embed each configuration in a two-dimensional space^37^.

#### Minimal pore radius

To estimate the minimal pore radius in the closed and open states we used HOLE^98^. The analysis was performed on 300 conformations taken with a stride of 1 ns from the last halves of the equilibration trajectories for each channel conformation, open and closed.

#### Water density along the central pore

From the equilibration and metadynamics trajectories, we extracted a set of frames using a stride of 1 ns. For every frame, we estimated the pore profile using HOLE^98^ and the three-dimensional map of water occupancy using the Volmap tool of VMD^99^. We integrated water occupancy in the xy plane (perpendicular to the central pore) and considered only the volume confined within the pore profile estimated by HOLE. For Figures 2B and S15 we used all the extracted frames, and for Figures 2A and S13 (average profiles) – the second halves of the trajectories.

#### Water density inside the peripheral cavities

To analyze the water density inside the PCs, we separated intracellular solution from the rest applying the following procedure. First, we calculated the 3D map of water occupancy using the Volmap tool of VMD^99^. We then generated a binary map by setting each voxel to either 0 or 1 depending on whether or not the local value of the density overcomes a predefined threshold. After that we used a clustering procedure to group the voxels using gmx cluster program of Gromacs 5.0.4 toolkit^94^. This approach allowed us to identify the cluster containing the intracellular solution and all the regions in space connected to it. Finally, we integrated the water occupancy of the selected cluster in the XY plane (perpendicular to the central axis).

#### Accessibility of S6 residues to the intracellular solution

To estimate the accessibility of the S6 residues to the intracellular solution, we considered the second halves of the equilibration trajectories. We applied the procedure described above to separate the coordinates of the intracellular water from the rest of the system. Finally, as a measure of accessibility, we calculated the overlap between the three-dimensional occupancy map of the intracellular water and that of each S6 residue.

#### Solvent accessible surface area (SASA)

Since we were only interested in the surface area accessible to the water contained the central pore and not in the PCs, we first defined the surface of the pore and then calculated the area pertaining to each S6 residue. In particular, we generated a binary three-dimensional map of the protein atoms (by setting the value of each voxel to either 0 or 1 depending on whether or not the local density exceeded a predefined threshold). We then extracted voxels with a value of 1 that were 1.5 Å away from voxels with value 0 (by doing this, we defined the surface as the region accessible to a sphere of radius 1.5 Å). We then calculated the occupancy map of each S6 residue and its overlap with the surface region. The overlap was re-scaled by dividing the raw value by the maximum surface area of the residue.

#### Analysis of free energy

The free energy profiles for ion permeation and for the closed-to-open transition were calculated using the reweighting approach introduced by Tiwary and Parrinello^45^. In the case of the ion permeation, the statistics from all non-biased ions was taken into account together with the biased ones to generate the free energy profile. The first parts of all metadynamics trajectories were not considered in the histograms. In particular, for the ionic transport in the closed and open states, the first 350 and 300 ns were ignored, respectively, while for the closed-to-open transition, the first 120 ns were ignored for all the temperatures. Free energy profiles were calculated at regular intervals and convergence of the simulations was ascertained by observing that, after a complete exploration of the CV space with several back and forth transition, the profiles did not change over time (Figures S3 and S6). Errors were calculated using the blocking transformation analysis^100^.

### Evolutionary Analysis

To investigate the possible evolutionary conservation of the peculiar profile of hydrophobicity shown by the S6 residues in rat TRPV1, we analyzed a comprehensive multiple sequence alignment encompassing 2851 members of the TRP channels family that we obtained previously^57^. To calculate, for each position of S6, the average solvation free energy and the corresponding standard deviation, we used the hydrophobic scale developed by Hessa et al.^42^.

### Experimental testing of a single residue mutant

#### Xenopus oocyte preparation and electrophysiology

Oocytes from female *Xenopus laevis* frogs were prepared using collagenase digestion, as describe earlier^101^. Briefly: oocytes were digested using 0.2 mg/ml collagenase (Sigma) in a solution containing

82.5 mM NaCl, 2 mM KCl, 1 mM MgCl_2_, and 5 mM HEPES, pH 7.4 (OR2) overnight for ∼ 16 h at 18 °C in a temperature controlled incubator. Oocytes were selected and kept in OR2 solution supplemented with 1.8 mM CaCl_2_ and 1% penicillin/streptomycin (Mediatech) at 18 °C. Point mutants of TRPV1 in the pGEMS oocyte vector were generated using the QuikChange kit (Agilent Technologies). cRNA was generated from linearized cDNA using the mMessage mMachine kit (Ambion). cRNA (50 ng) was microinjected into each oocyte using a nanoliter injector system (World Precision Instruments). The experiments were performed 5 days after injection.

Two-electrode voltage clamp (TEVC) measurements were performed as described earlier^88^ in a solution containing 97 mM NaCl, 2 mM KCl, 1 mM MgCl_2_, 5 mM 2-(N-Morpholino)ethanesulfonic acid (MES) and 5 mM HEPES, pH 7.4. MES was added to our usual extracellular oocyte buffer to be able set the to pH to 4 with HCl in the stimulating solution. Currents were recorded with thin-wall inner filament-containing glass pipettes (World Precision Instruments) filled with 3 M KCl in 1% agarose. Currents were measured with a ramp protocol from -100 to 100 mV performed once 0.5 second from a holding potential of 0 mV, and the currents at -100, 0 and 100 mV were plotted.

#### Cell culture and Ca^2+^ imaging

Human embryonic kidney 293 (HEK293) cells were obtained from the ATCC and were cultured in minimal essential medium (MEM) (Invitrogen) containing supplements of 10% (v/v) Hyclone-characterized Fetal Bovine Serum (FBS) (Thermo Scientific), 100 IU/ml penicillin, and 100 μg/ml streptomycin. Transient transfection was performed at ∼ 70% cell confluence with the Effectene reagent (QIAGEN) according to the manufacturer’s protocol. Cells were incubated with the lipid-DNA complexes overnight. Then, 24 h post-transfection, cells were trypsinized and replated on poly-l-lysine-coated glass coverslips and incubated for an additional 24 h (in the absence of the transfection reagent) before measurements. Cultured HEK cells were kept in a humidity-controlled tissue-culture incubator maintaining 5% CO_2_ at 37°C.

Ca^2+^ imaging measurements were performed with an Olympus IX-51 inverted microscope equipped with a DeltaRAM excitation light source (Photon Technology International). HEK cells were loaded with 2 μM low affinity Fura-2 AM (Abcam) at 37°C in MEM – FBS for 45 min before the measurement, and dual-excitation images at 340 and 380 nm were recorded with a Roper Cool-Snap digital CCD camera. The standard extracellular solution used in all experiments contained (in mM): 137 NaCl, 5 KCl, 1 MgCl_2_, 2 CaCl_2_, 10 HEPES and 10 glucose, pH 7.4 (adjusted with NaOH). For the low pH solution the composition was modified to 137 NaCl, 5 KCl, 1 MgCl_2_, 2 CaCl_2_, 5 HEPES, 5 MES and 10 glucose, pH 4.0 (adjusted with HCl). Low pH and capsaicin solution (stock in EtOH) were applied with a gravity driven whole chamber perfusion system. Heat stimulation was performed with custom made system, and solution temperature was continuously recorded with a thermo probe located in a close proximity to the recording field. Data analysis was performed using the Image Master software (Photon Technology International).

